# Influenza A Virus Disruption of Dendritic Cell-Natural Killer Cell Crosstalk Impacts Activation of Helper and Cytotoxic T cell Subsets

**DOI:** 10.1101/2024.02.01.578467

**Authors:** Connor A. Morson, Chandana K. Uppalapati, Brina S. Lopez, Lisa M. Kronstad

## Abstract

Dendritic cells (DC) and Natural killer (NK) cells engage in reciprocal interactions to trigger an efficient innate immune response while governing the adaptive immune response. Here we used an *ex vivo* autologous human primary immune cell co-culture of DCs and NK cells to investigate the impact of DC-NK cell crosstalk on activation of CD4^+^ and CD8^+^ naïve T cell responses to influenza A viral (IAV) infection. Using multiparameter flow cytometry, we observed that culturing T cells with DC and NK cells led to enhanced expression of CD69 and CD25 activation markers and increased proliferative ability of both CD4^+^ and CD8^+^ T cell subsets. Exposure of DCs to the pandemic A/California/07/2009 (H1N1) strain in NK cell co-culture led to a reduced frequency of CD4^+^CD69^+^, CD8^+^CD69^+^, CD4^+^CD25^+^, CD8^+^CD25^+^ T cell subsets and a reduced expansion of CD4^+^ T cells. The IAV-mediated curtailment of T cell activation was dependent on the ability of A/California/07/2009 (H1N1) to replicate as inactivation of the virus rescued expression of CD69, CD25 on both CD4^+^ and CD8^+^ T cell subsets and triggered expansion of CD4^+^ T cells. Further, we discovered exposure of DCs to the A/Victoria/361/2011 (H3N2) IAV strain also significantly impaired expression of CD69 on CD4^+^ and CD8^+^ T cells and CD25 on CD8^+^ T cells. In contrast with the A/California/07/2009 (H1N1 strain), inactivation of A/Victoria/361/2011 (H3N2) failed to fully restore T cell expression of CD69 and CD25 and proliferation. Collectively, these data demonstrate that IAV partially usurps the ability of DC-NK cell crosstalk to activate naïve CD4^+^ and CD8^+^ T cells in a strain-dependent manner. These data may inform the immunological signals required to trigger a potent cellular immune response to IAV, which may elicit broader and more durable protection than current inactivated vaccine platforms.

## Introduction

The World Health Organization estimates the annual global burden of seasonal influenza at 1 billion cases, including 3-5 million severe infections and 290,000-650,000 deaths [1]. Current influenza vaccines elicit antibodies that predominantly target the highly variable head region of hemagglutinin and effectiveness is limited by antigenic drift and failure to induce bone marrow-residing long-lived plasma cells [2–5]. Vaccines that confer durable protection while broadly protecting against seasonal drift variants and against potential pandemic strains are urgently needed [6–8]. Prior work has shown that influenza-specific T cells elicit broad viral strain coverage and provide protective immunity in mice, non-human primates, and humans—likely due to the ability to recognize epitopes derived from highly conserved internal viral proteins, such as nucleoprotein, that poorly tolerate amino acid changes [9–13]. Vaccination with mRNA packaged in lipid nanoparticles (LNPs) allows uptake by dendritic cells (DCs) and subsequent induction of potent T cell responses–rendering this vaccine platform attractive for the next generation of influenza vaccines [14].

Initiating antigen-specific CD8^+^ T cell responses relies on DCs, which are equipped with pattern recognition receptors to sense incoming pathogens and possess the unique ability to prime both naïve CD4^+^ and CD8^+^ T cells by presenting and cross-presenting influenza A virus (IAV) antigen [15]. DCs influence helper T cell differentiation by polarizing CD4^+^ helper T cells toward T_H_1 or T_H_2 response based on the environment in which the DC has been stimulated [16]. DC function is tightly regulated by interactions with natural killer (NK) cells, with NK cells acting as a rheostat capable of influencing the magnitude and type of DC-mediated T cell activation [17–21]. Indeed, experiments performed with IAV-infected mice showed that NK cells were required for DC migration and CD8^+^ T cell priming in an interferon-γ (IFN-γ)-dependent mechanism [22]. Further, the ability of NK cells to lyse virally-infected cells leads to the generation of viral antigenic material; viral debris is then engulfed by DCs and viral antigens are processed and presented by mature DCs to both CD8^+^ (cross-priming) and CD4^+^ T cells [23].

Our prior data established that DC-NK cell crosstalk leads to heightened expression of T cell surface activation markers including CD69 and CD25 on CD3^+^ T cells–comprising both CD4^+^ helper and CD8^+^ cytotoxic subsets. This heightened T cell activation profile was curtailed upon when DCs were exposed to multiple IAV strains including A/California/07/2009 (H1N1) (Cal/09) or the A/Victoria/361/2011 (H3N2) (Vic/11). However, the relative impact of IAV on DC-NK cell crosstalk on CD4^+^ helper and CD8^+^ cytotoxic T cell unique subsets including surface activation and their subsequent proliferative ability remained to be established.

To address these questions, we used an *ex vivo* autologous human cell triple co-culture system to establish the role of DC-NK cell crosstalk on the relative activation and proliferation of CD4^+^ helper and CD8^+^ cytotoxic T cells in response to Cal/09 and Vic/11. Further, we aimed to test the hypothesis that active viral protein synthesis is required to curtail T cell activation by using replication-defective virus. These viral strains were selected based on our studies that demonstrated that NK cells secrete higher levels of IFN-γ response to Cal/09 compared to Vic/11, and this translates into heightened T cell IFN-γ production after DC-NK cell crosstalk [24,25]. We found that the DC-NK cell crosstalk heightened the expression of early (CD69) and late (CD25) markers of T cell activation while triggering the proliferation of both subsets. Exposure of DCs to IAV impaired both surface activation and proliferation in a strain- and replication-dependent manner, supporting a role for active viral protein synthesis in abrogating T cell activation.

## Materials and Methods

### Virus Production and Titration

A/Victoria/361/2011 (Vic/11; H3N2) (BEI Resources) and A/California/07/2009 (Cal/09: H1N1) (BEI Resources) viruses were propagated by inoculation from a seed stock in 10-day-old specific pathogen-free embryonated chicken eggs (Charles River Laboratories Wilmington, MA) at 35°C with 55% - 65% humidity. Infection was allowed to proceed for 48 hours (h), followed by a 16 h incubation at 4°C and then harvesting of the allantoic fluid. Aliquots of virus stocks were prepared and stored at -80°C. Each batch of the infectious Vic/11 and Cal/09 virus titer was determined by a standard plaque assay on MDCK cells in the presence of 2 µg/mL L-1-tosylamide-2-phenylethyl chloromethyl ketone (TPCK)-treated trypsin. Replication-defective viruses were prepared by delivering 2400 µJ/cm2 of UV light (254 nm) using a Strata linker 1800 (Stratagene, La Jolla, CA). Inactivation of Cal/09 (Cal09-UV) and of Vic/11 (Vic/11-UV) were verified both by MDCK plaque assay and by intracellular staining with influenza nucleoprotein antibody followed by flow cytometry.

### Peripheral Blood Mononuclear Cell Isolation

Human peripheral blood mononuclear cells (PBMCs) were isolated from whole blood collected from healthy adult donors as previously described [25]. Briefly, antecubital venipuncture was performed on both male and female donors and whole blood was collected into BD Vacutainer^TM^ with Sodium Heparin blood collection tubes (BD, Franklin Lakes, NJ). Immediately following collection, blood was diluted 2-fold in calcium- and magnesium-free phosphate-buffered saline (PBS pH: 7.2; Fisher Scientific, Waltham, MA). PBMCs were isolated from whole blood using density gradient centrifugation at 1,200 x *g* for 10 min where diluted blood was layered over 15 mL of Lymphoprep™ (STEMCELL Technologies, Vancouver, Canada) followed by harvesting of the buffy coat. Residual red blood cells were lysed by 5 mL of ACK lysing buffer (Gibco, Waltham, MA) for 10 mins. PBMCs were washed twice with 1X PBS, pelleted, suspended and cryopreserved at -80°C in 90% (v/v) fetal bovine serum (FBS; R&D Systems, Minneapolis, MN) and 10% (v/v) dimethyl sulfoxide (DMSO; Corning, Corning, NY). PBMC vials were labeled with the unique alphanumerical code assigned to each donor. All samples were transferred to liquid nitrogen storage within 24 h.

### Human Monocyte-derived Dendritic cell (MoDCs) Generation

Human monocyte-derived dendritic cells (MoDCs) were generated as previously described [26]. Briefly, monocytes were isolated from cryopreserved PBMCs using magnetic activated cell sorting (MACS®), and Pan Monocyte Isolation kit (Miltenyi Biotec, Auburn, CA, CAT#130-096-537) following the manufacturer’s protocol. Monocytes were enriched by plating in serum-free medium in 12-well Nunclon™ Delta surface-treated flat bottom plates (Thermo Fisher Scientific, Waltham, MA) at a density of 0.5 - 1.0 x 10^6^ cells/well followed by incubation at 37°C with 5% CO_2_. 2 h post-incubation, adherent monocytes were cultured in RPMI 1640 medium supplemented with 10% FBS, 1% L-Glutamine, and 1% Penicillin/Streptomycin (RP10) (Thermo Fisher Scientific, Waltham, MA), recombinant human granulocyte-macrophage colony-stimulating factor (rhGM-CSF) (300U/mL; Thermo Fisher Scientific, Waltham, MA) and recombinant human interleukin 4 (rhIL-4) (300 U/mL; Thermo Fisher Scientific, Waltham, MA). On day 2 and day 5 post-monocyte isolation, half of the medium volume was supplemented with fresh medium containing cytokines. MoDC differentation was confirmed by analytical flow cytometry measuring the downregulation of a monocyte surface marker (CD14) and upregulation of a DC surface marker (CD209) [25].

### Cell purification, Stimulation, and Infection

PBMCs were thawed and washed in RP10 medium (RPMI 1640 medium supplemented with 10% FBS plus 2 mM L-glutamine, 100 U/ml penicillin, and 100 U/ml streptomycin). Cell viability as determined using a 0.4% Trypan Blue solution (Sigma) in a hemocytometer. MoDCs were removed from 12-well Nunclon™ Delta surface-treated flat bottom plates (Thermo Fisher Scientific, Waltham, MA), pelleted, counted and re-suspended in serum-free RPMI. MoDCs were then plated into wells of a 96-well U-bottom tissue culture-treated plate (CELLTREAT, Pepperell, MA) at a seeding density of 1 x 10^6^ cells per well. MoDCs were exposed to either Cal/09 or Vic/11 at a multiplicity of infection (MOI) of 3 for 1 h at 37°C and 5% CO_2_. Viral inoculum in serum-free media or serum-free media alone (mock condition) was removed at 1 HPI via centrifugation and cells resuspended in 200 μl RP10. The Human NK Cell Isolation Kit (Miltenyi Biotec; CAT#130-092-657) was used to purify NK cells and the Human Naïve Pan T Cell Isolation kit (Miltenyi Biotec; CAT#130-097-095) was used to purify T cells. The Naïve Pan T Cell kit yields cells that are CD3^+^CD45RA^+^CD197^+^ by using specific antibodies to CD45RO, CD14, CD15, CD16, CD19, CD25, CD34, CD36, CD57, CD123, HLA-DR, CD235a and CD244. NK cells and pan naïve T cells were co-cultured with MoDCs in serum-containing RP10 medium at a seeding density of 4 x 10^6^ and 1 x 10^6^ cells per well, respectively. 10 U/ml of IL-2 was added to each well containing naïve T cells [27]. Anti-CD3/CD28-coated Dynabeads^®^ (Gibco, Life Technologies; CAT#11161D) were used as a positive control for antigen-independent stimulation following the manufacturer’s instructions at a 1:1 bead-to-cell ratio at the indicated time points.

### Staining and Flow Cytometry

After magnetic isolation, cell purity was assessed using analytical flow cytometry. To determine the purity of MoDCs, cells were stained with APC/Cyanine7 anti-human CD14 (clone: 63D3 BioLegend®) and Brilliant Violet 421™ anti-human CD209 (clone: 9E9A8 BioLegend®) antibodies. For MoDC, NK cell, and T cell co-culture activation experiments, cells were first stained with Live-or-Dye™ 405/545 (Biotium, Fremont, CA) followed by staining with PE anti-human CD69 (clone: FN50; BioLegend®), or PE anti-human CD25 (clone: BC96; BioLegend®) antibodies. For proliferation experiments, T cells were stained with carboxyfluorescein diacetate succinimidyl ester (CFSE) (eBioscience/Thermo Fisher Scientific, Waltham, MA) at the time of co-culture on day 7 followed by Live-or-Dye™ 405/545 viability stain (Biotium, Fremont, CA) on day 13. Cells were fixed with 1X BD FACS™ Lysing Solution (BD Biosciences) and permeabilized with 1X FACS Perm II (BD Biosciences) buffers following the manufacturer’s protocol. For intracellular staining, cells were stained with Pacific Blue™ anti-human CD3 (clone: UCHT1; BioLegend®), Alexa Fluor 647 anti-human CD4 (clone: RPA-T4; BioLegend®), and PE-Cyanine7 anti-human CD8 (Clone: SK1; BioLegend®) antibodies. Uncompensated data was collected by performing analytical flow cytometry (Guava® easyCyteTM 12HT flow cytometer (Luminex Corporation, Austin, TX)). Data was compensated and analysed using FlowJo™ version 10.6.1 (BD Biosciences). Fluorescence Minus One (FMO) was used as gating control and was used in conjunction with its corresponding compensation to allow for the gating of the positive population. Gating schematics for each cell population are shown in **Figure S1**.

### Statistical Analysis

Paired Wilcoxon signed-rank tests were performed using R. An alpha value of 0.05 was set to all data analysis and a p-value of < 0.05 was considered statistically significant.

### IRB Approval-Ethics

The research protocol for this study was approved by Midwestern University’s Institutional Review Board (IRB) under study number IRB/AZ 1299. This study involved male and female adult human donors self-identifying as healthy, between the ages of 18-65.

## Results

### MoDC-NK cell crosstalk increases the frequency of CD69^+^CD4^+^ and CD69^+^CD8^+^ T cells

In our prior study, we found that DC-NK cell crosstalk triggered an increase in CD69^+^ expression on pan-naïve T cells [25]. The appearance of CD69 on the plasma membrane is a classic early marker of lymphocyte activation and has multifaceted roles including regulation of cytokine release and homing and migration of lymphocytes [28]. Therefore, we asked whether DC-NK crosstalk influences CD69 expression preferentially on helper CD4^+^, on cytotoxic CD8^+^, or on both subsets [25]. We used a co-culture system consisting of monocyte-derived dendritic cells (MoDCs) either mock-treated or exposed to A/California/07/2009 pandemic H1N1 (Cal/09) strain or the A/Victoria/361/2011 H3N2 (Vic/11) at an MOI of 3 with autologous NK cells and naïve T cells for 47 hours (h). T cells were then evaluated for changes in CD69 expression by analytical flow cytometry [29]. Anti-CD3/28-coated Dynabeads^®^ were used as a positive control for antigen-independent stimulation, which indeed triggered CD69 expression on both helper CD4^+^ and cytotoxic CD8^+^ subsets **(Figure 1A**). We first asked whether naïve T cell activation was influenced by the presence of MoDCs and/or NK cells. To this end, T cells were cultured either alone (0:0:1), with MoDCs (1:0:1), with NK cells (0:4:1), or in triple co-culture at a MoDC: NK cell: T cell ratio of 1:4:1. This cell ratio was found to robustly activate pan naïve T cells in a prior study [25]. We found that both helper CD4^+^ (median = 62.5%) and cytotoxic CD8^+^ (median = 57.2%) T cell subsets exhibited significantly lower frequencies of the CD69^+^ population when co-cultured with MoDCs and NK cells, compared to only MoDCs, only NK cells, or solely T cells **(Figure 1A-B**).

**Figure 1.**
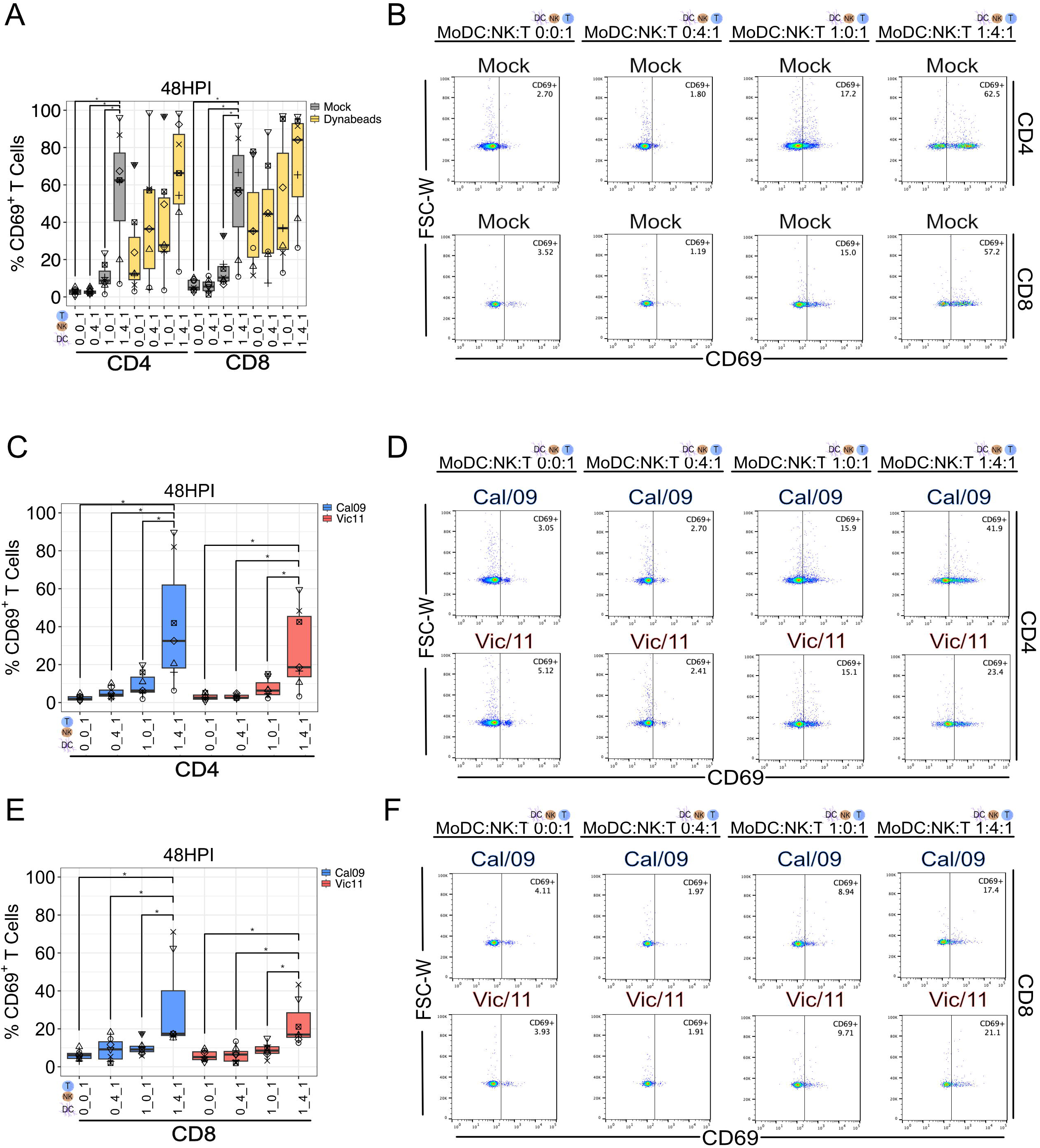
MoDCs trigger expression of CD69 on CD4^+^ and CD8^+^ T cells in a NK cell-dependent manner. **(A)** Summary plot of CD69^+^ frequency on CD4^+^ and CD8^+^ T cell subset cultured alone (0:0:1) for 48 hours (h) or at the indicated MoDC:NK:T cell ratios of 0:4:1, 1:0:1, 1:4:1 for 47 h under after mock treatment. Anti-CD3/28-coated Dynabead^®^ treatment at a 1:1 bead-to-cell ratio for the last 6 h of the co-culture served as a positive control (*n* = 7). **(B)** Representative flow plots showing the frequency of CD4^+^CD69^+^ (top panel) and CD8^+^CD69^+^ (bottom panel) T cell subsets after 47 h co-culture with mock-treated MoDCs and NK cells at the indicated cell ratio. Summary plot **(C)** and representative **(D)** flow plots of CD69^+^ frequency on CD4^+^ T cells cultured alone (0:0:1) for 48 h or at the indicated MoDC:NK:T cell ratios of 0:4:1, 1:0:1, 1:4:1 for 47 h under after MoDC exposure to Cal/09 or Vic/11 (MOI = 3) (*n* = 7). Summary plot **(E)** and representative **(F)** flow plots of CD69^+^ frequency on CD8^+^ T cells cultured alone (0:0:1) for 48 h or at the indicated MoDC:NK:T cell ratios of 0:4:1, 1:0:1, 1:4:1 for 47 h under after MoDC exposure to Cal/09 or Vic/11 (MOI = 3) (*n* = 7). The data point with the same shape between conditions indicates that it is derived from the same donor. *p < 0.05, Wilcoxon signed-rank test.

We next sought to evaluate how influenza impacts MoDC-NK crosstalk and subsequent T cell activation. MoDCs were exposed to either Cal/09 or Vic/11 prior to co-culture with autologous NK and naïve T cells. Similar to mock conditions, we found that the frequency of helper CD4^+^ (**Figure 1C-D**) and cytotoxic CD8^+^ (**Figure 1E-F**) subsets was highest when the T cells were in cultured with MoDCs and NK cells (1:4:1) compared to when cultured independently (0:0:1 with only MoDCs (1:0:1), or only NK cells (0:4:1). This pattern was observed when MoDCs were exposed to either the Cal/09 or Vic/11 viral strains (**Figure 1C-F**). Collectively, these data support a role for MoDC-NK cell crosstalk in stimulating T cells, as assessed by expression of CD69, under either steady-state or IAV exposure conditions.

### IAV impairs MoDC-NK cell-mediated increase in the frequency of CD69 expression of CD4^+^ and CD8^+^ T cells in a replication-dependent manner

Next, we directly evaluated whether IAV exposure had an impact on CD4^+^ and CD8^+^ T cell activation, compared to the mock, uninfected condition. We found that exposure of MoDCs to either Cal/09 or Vic/11 yielded significantly lower levels of CD4^+^CD69^+^ (Cal/09: median = 32.5%; Vic/11: median = 18.6%) and CD8^+^CD69^+^ (Cal/09: median = 17.4%; Vic/11: median = 17%) T cells compared to cells cultured with mock-treated MoDCs when in triple co-culture (**Figure 2A-B**). We next asked whether this lower level of CD69 expression was dependent on viral replication by rendering the virus replication-defective using ultraviolet (UV) light, which forms dimers between consecutive nucleotides and interferes with genomic replication and transcription [30,31]. When MoDCs were exposed to UV-inactivated, replication-defective virus, T cell CD69^+^ expression was significantly increased by Cal/09-UV in both subsets (CD4: median of RC = 32.5%, median of UV = 56.3%; CD8: median of RC = 17.4%, median of UV = 52.2%) (**Figure 2C-D**). While the median values were also higher after exposure to Vic/11-UV compared to replication-competent (Vic/11-RC) virus, the increase was not statistically significant on either the CD4^+^ (RC median = 18.6, UV media = 37.5) or the CD8^+^ subset (RC median = 17, UV median = 30) (**Figure 2C-D**). Collectively, these data support a model where early CD4^+^ and CD8^+^ T cell activation is heightened by MoDC-NK cell crosstalk and that this activation is impaired when MoDCs are exposed to either the Cal/09 or Vic/11 IAV strains. Further, T cell CD69 expression is restored when Cal/09 is rendered replication-defective–supporting a role for active viral protein synthesis by Cal/09 in abrogating T cell activation under these conditions.

**Figure 2.**
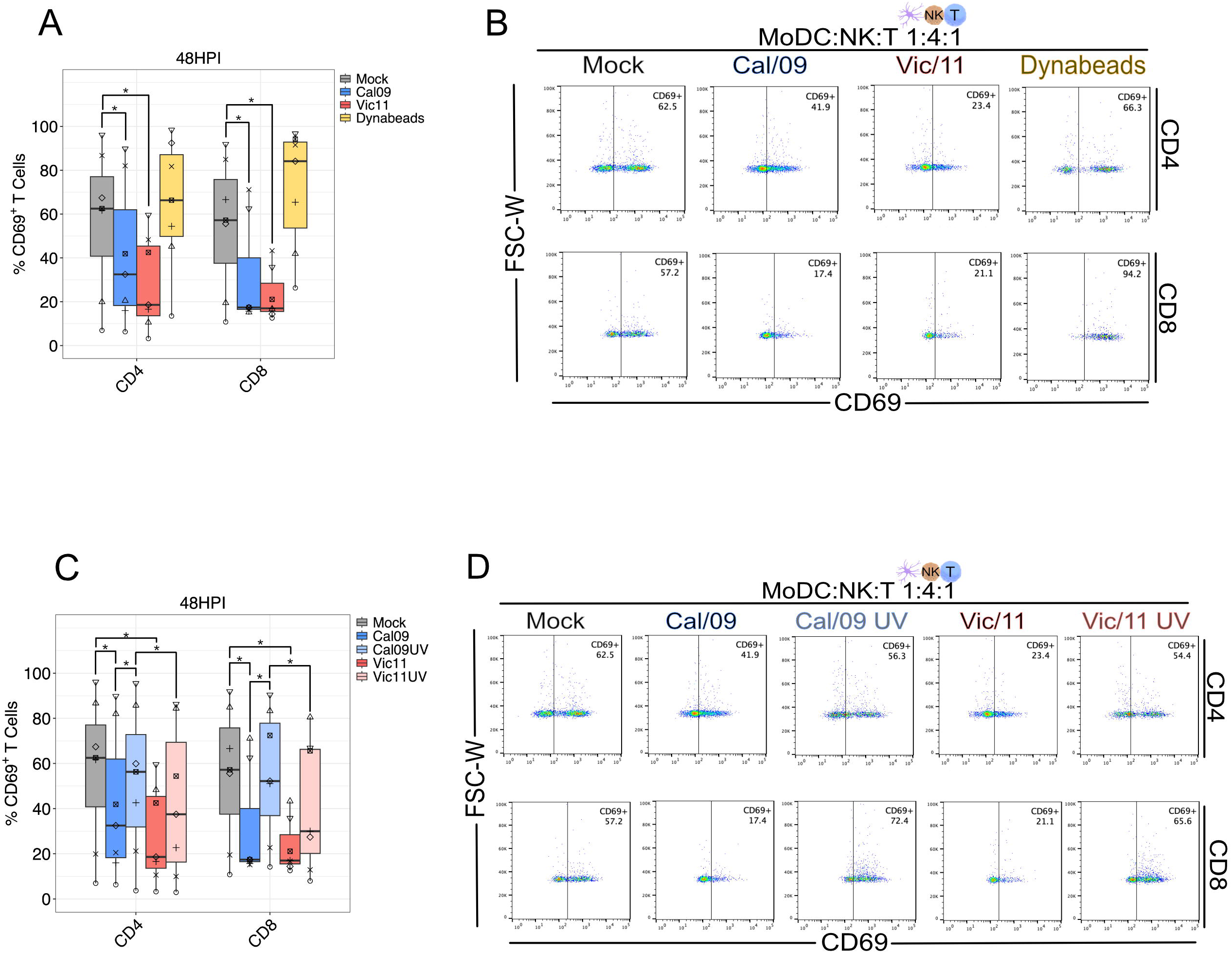
Replication of IAV is required for impairment of CD4^+^ and CD8^+^ T cell expression of CD69 after MoDC-NK cell co-culture. **(A)** Summary plot of CD69^+^ frequency on CD4^+^ and CD8^+^ T cells after 47 h co-culture with mock-treated or Cal/09 or Vic/11-exposed MoDCs (MOI = 3) and NK cells at MoDC:NK:T cell ratio of 1:4:1 (*n* = 7). **(B)** Representative flow plots showing the frequency of CD69^+^ on CD4^+^ (top panel) and CD8^+^ (bottom panel) T cell subsets after 47 h co-culture with mock-treated or Cal/09 or Vic/11-exposed MoDC (MOI = 3) and NK cells at MoDC:NK:T cell ratio of 1:4:1. **(C)** Summary plot of CD69^+^ frequency on CD4^+^ and CD8^+^ T cell subsets after 47 h co-culture with mock-treated, replication-competent or UV-inactivated viral-exposed MoDC (MOI = 3) and NK cells at MoDC:NK:T cell ratio of 1:4:1 (*n* = 7). **(D)** Representative flow plots showing the frequency of CD69^+^ on CD4^+^ (top panel) and CD8^+^ (bottom panel) T cell subset after 47 h co-culture with mock-treated, replication-competent or UV-inactivated Cal/09 and Vic/11 (MOI = 3) and NK cells at MoDC:NK:T cell ratio of 1:4:1. Anti-CD3/28-coated Dynabead^®^ treatment at a 1:1 bead-to-cell ratio for the last 6 h of the co-culture served as a positive control. The data point with the same shape between conditions indicates that it is derived from the same donor. *p < 0.05, Wilcoxon signed-rank test.

### MoDC-NK cell crosstalk increases the frequency of CD25^+^CD4^+^ and CD25^+^CD8^+^ T cells

Next, we evaluated the impact of MoDC-NK cell crosstalk and IAV on the expression of CD25 (IL-2 receptor alpha chain), on the surface of helper CD4^+^ and cytotoxic CD8^+^ T cells, using the same experimental design that was used to evaluate CD69 expression. The relative expression of CD25 on the subsets is of interest as previous studies of viral infection in mice found that induction of CD25 expression on antigen-specific CD8^+^ T cells was dependent on the presence of CD4^+^ T cells [32]. CD25 was evaluated at 96 HPI based on prior investigation of CD25 expression kinetics on naïve human T cells [33]. We used T cells exposed to anti-CD3/CD28-coated Dynabeads^®^ for 21 h as a positive control for antigen-independent stimulation (**Figure 3A**). We evaluated whether our data recapitulated the earlier finding that pan CD3^+^ T cell–comprised of both CD4^+^ and CD8^+^ T cell subsets–had the highest percentage of CD25^+^ in the triple co-culture condition that included both MoDCs and NK cells, or if one of the subsets was affected by MoDC-NK cell crosstalk at a greater extent than the other [25]. Indeed, we found that both CD4^+^ and CD8^+^ T cells exhibited increased CD25^+^ expression when co-cultured with MoDCs and NK cells, compared to only MoDCs, only NK cells, or solely T cells **(Figure 3A-B)**.

**Figure 3.**
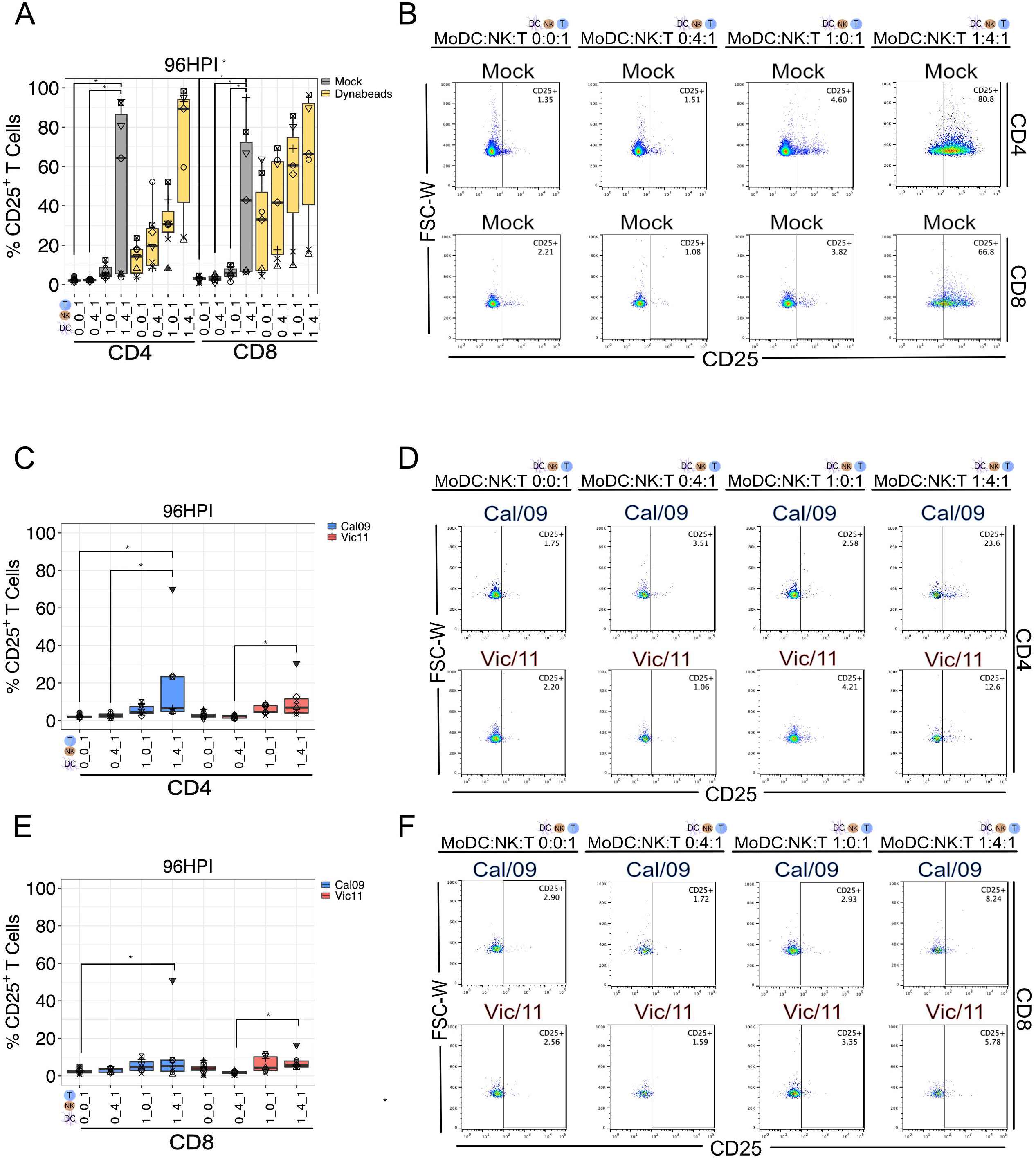
MoDCs trigger expression of CD25 on CD4^+^ and CD8^+^ T cells in a NK cell-dependent manner. **(A)** Summary plot of CD25^+^ frequency on CD4^+^ and CD8^+^ T cell subsets cultured alone (0:0:1) for 96 h or at the indicated MoDC:NK:T cell ratios of 0:4:1, 1:0:1, 1:4:1 for 95 h after mock treatment (*n* = 7). Anti-CD3/28-coated Dynabead^®^ treatment at a 1:1 bead-to-cell ratio for the last 21 h of the co-culture served as a positive control. **(B)** Representative flow plots showing the frequency of CD25^+^ on CD4^+^ (top panel) and CD8^+^ (bottom right panel) T cell subsets after 95 h co-culture with mock-treated MoDCs and NK cells at the indicated cell ratio. Summary plot **(C)** and representative **(D)** flow plots of CD25^+^ frequency on CD4^+^ T cells cultured alone (0:0:1) for 96 h or at the indicated MoDC:NK: T cell ratios of 0:4:1, 1:0:1, 1:4:1 for 95 h under after MoDC exposure to Cal/09 or Vic/11 (MOI = 3) (*n* = 7). Summary plot **(E)** and representative **(F)** flow plots of CD25^+^ frequency on CD8^+^ T cells cultured alone (0:0:1) for 96 h or at the indicated MoDC:NK:T cell ratios of 0:4:1, 1:0:1, 1:4:1 for 95 h under after MoDC exposure to Cal/09 or Vic/11 (MOI = 3) (*n* = 7). The data point with the same shape between conditions indicates that it is derived from the same donor. *p < 0.05, Wilcoxon signed-rank test.

Next, we evaluated the impact of MoDC-NK crosstalk on T cell CD25 expression after exposing the MoDCs to Cal/09 or Vic/11, prior to co-culture with autologous NK and naïve T cells. In contrast with the increased activation from MoDC-NK cell crosstalk observed under steady-state conditions, we found no significant difference in the frequency of CD4^+^CD25^+^ (**Figure 3C-D**) and CD8^+^CD25^+^ (**Figure 3E-F**) populations between the triple co-culture condition (1:4:1) and the MoDC and T cell co-culture (1:0:1). Notably, after MoDC exposure to Cal/09 or Vic/11, the frequency of CD4^+^CD25^+^ in triple co-culture (1:4:1) was significantly higher than with only NK and T cells (0:4:1). After MoDC exposure to Cal/09 only, but not to Vic/11, the frequency of CD4^+^CD25^+^ in triple co-culture (1:4:1) was significantly higher than with only T cells (0:0:1). Together, these data show that culturing T cells with both MoDCs and NK cells leads to heightened expression of CD25 (IL-2Rα) on both helper CD4^+^ and cytotoxic CD8^+^ naïve T cells, however this T cell activation is curtailed when MoDCs are exposed to IAV.

### IAV impairs MoDC-NK cell-mediated increase in the frequency of CD25 expression of CD4^+^ and CD8^+^ T cells in a replication-dependent manner

We next directly compared the effect of IAV on CD25^+^ expression for both T cell subsets when in triple co-culture (1:4:1). We found that when CD8^+^ T cells were exposed to NK cells and Cal/09- or Vic/11-exposed MoDCs they experienced a significant reduction in CD25^+^ expression when compared to the mock-treated MoDCs **(Figure 4A-B)**. We also observed that there was significantly increase in CD25 expression on CD4^+^ T cells than CD8^+^ T cells when exposed to Cal/09 virus **(Figure 4A-B)**. Additionally, both CD4^+^ and CD8^+^ T cells expressed significantly higher levels of CD25^+^ when exposed to Cal/09-UV vs replication-competent Cal/09 (CD4: RC median = 6.52%, UV median = 83.3%; CD8: RC median = 5.23%, UV median = 67%) **(Figure 4C-D)**. Lastly, CD25 expression was significantly higher on both CD4^+^ and CD8^+^ T cells when exposed to Cal/09-UV when compared to Vic/11-UV for both CD4^+^ (Cal/09-UV median = 83.3%, Vic/11-UV median = 15.6%) and CD8^+^ (Cal/09-UV median = 67%, Vic/11-UV median = 19.4%) T cells **(Figure 4C-D)**. Taken together, these data show that expression of the alpha chain of the IL-2 receptor (CD25) on helper CD4^+^ and cytotoxic CD8^+^ T cell activation is heightened by MoDC-NK cell crosstalk and this activation is significantly impaired when MoDCs are exposed to either the Cal/09 (CD4^+^ and CD8^+^ subsets) or Vic/11 (CD8^+^ subset) IAV strains. Further, active viral protein synthesis by Cal/09 is required for the observed impairment of T cell CD25 expression on both subsets.

**Figure 4.**
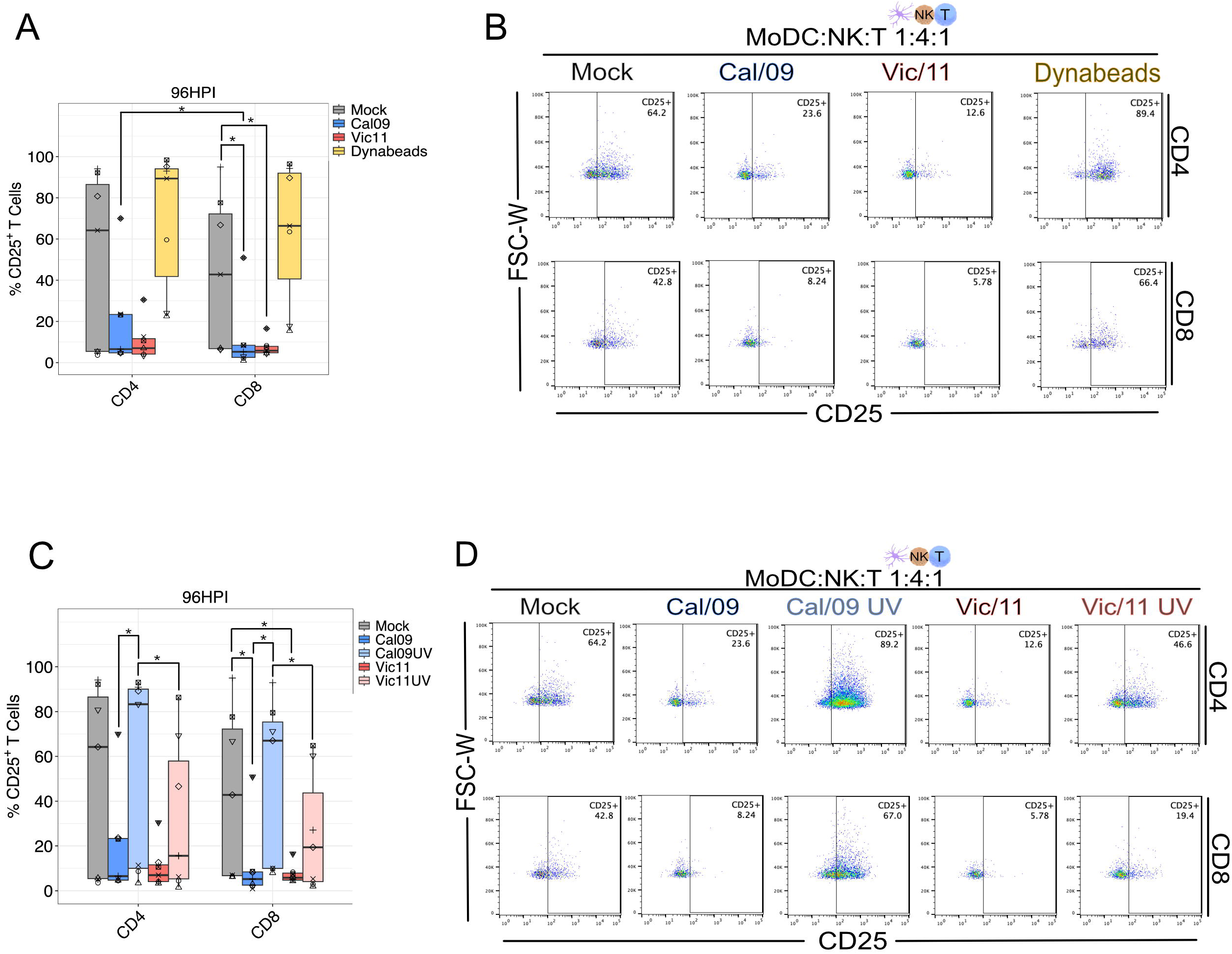
Replication of IAV is required for impairment of CD4^+^ and CD8^+^ T cell expression of CD25 after MoDC-NK cell co-culture. **(A)** Summary plot of CD25^+^ frequency on CD4^+^ and CD8^+^ T cells after 95 h co-culture with mock-treated or Cal/09 or Vic/11-exposed MoDCs (MOI = 3) and NK cells at MoDC:NK:T cell ratio of 1:4:1 (*n* = 7). **(B)** Representative flow plots showing the frequency of CD25^+^ on CD4^+^ (top panel) and CD8^+^ (bottom panel) T cell subsets after 95 h co-culture with mock-treated or Cal/09 or Vic/11-exposed MoDCs (MOI = 3) and NK cells at MoDC:NK:T cell ratio of 1:4:1. **(C)** Summary plot of CD25^+^ frequency on CD4^+^ and CD8^+^ T cell subsets after 95 h co-culture with mock-treated, replication-competent or UV-inactivated viral-exposed MoDCs (MOI = 3) and NK cells at MoDC:NK:T cell ratio of 1:4:1 (*n* = 7). **(D)** Representative flow plots showing frequency of CD25^+^ on CD4^+^ (top panel) and CD8^+^ (bottom panel) T cell subset after 95 h co-culture with mock-treated, replication-competent or UV-inactivated Cal/09 and Vic/11 (MOI = 3) and NK cells at MoDC:NK:T cell ratio of 1:4:1. Anti-CD3/28-coated Dynabead^®^ treatment at a 1:1 bead-to-cell ratio for the last 21 h of the co-culture served as a positive control. The data point with the same shape between conditions indicates that it is derived from the same donor. *p < 0.05, Wilcoxon signed-rank test.

### MoDC-NK cell crosstalk triggers the proliferation of CD4^+^ and CD8^+^ naïve T cells

Given the reduced frequencies of CD4^+^CD25^+^ and CD8^+^CD25^+^ subsets when MoDC were exposed to IAV and the role of IL-2 in T cell expansion, we next evaluated CD4^+^ and CD8^+^ T cell proliferation by measuring the intracellular dilution of CFSE-labeled T cells over a period of time, which permits permitting tracking of 8-10 cycles of cell proliferation. T cells exposed to anti-CD3/CD28-coated Dynabeads^®^ for 144 h served as the positive control for antigen-independent stimulation for the length of the co-culture (**Figure 5A**). Using the expansion index–a value that reflects the extent of how much the cell population has grown by dividing the total number of cells by the number of cells at the start of the culture–we found both CD4^+^ and CD8^+^ T cells underwent more cycles of cell division when co-cultured with MoDCs and NK cells (1:4:1), compared to with only MoDCs (1:0:1), with only NK cells (0:4:1), or independently (0:0:1) **(Figure 5A-B)**. The same pattern was observed for CD4^+^ **(Figure 5C-D)** and CD8^+^ **(Figure 5E-F)** T cells upon MoDC exposure to the Cal/09 IAV strain. For Vic/11, increased expansion of CD4^+^ T cells was observed in the triple co-culture (1:4:1) compared to when cultured solely with NK cells (0:4:1) or independently (0:0:1) but not when compared to co-culture solely with MoDCs (1:0:1). When MoDCs were exposed to Vic/11, CD8^+^ T cells did not expand in either the triple (1:4:1) or co-cultures (1:0:1; 0:4:1) although expansion was detected when compared to T cells cultured in isolation (0:0:1). These data support that MoDC-NK cell crosstalk is not only vital for CD4^+^ and CD8^+^ T cell subset activation of surface markers but for proliferation under steady-state conditions and when MoDCs are exposed to Cal/09, but not to Vic/11.

**Figure 5.**
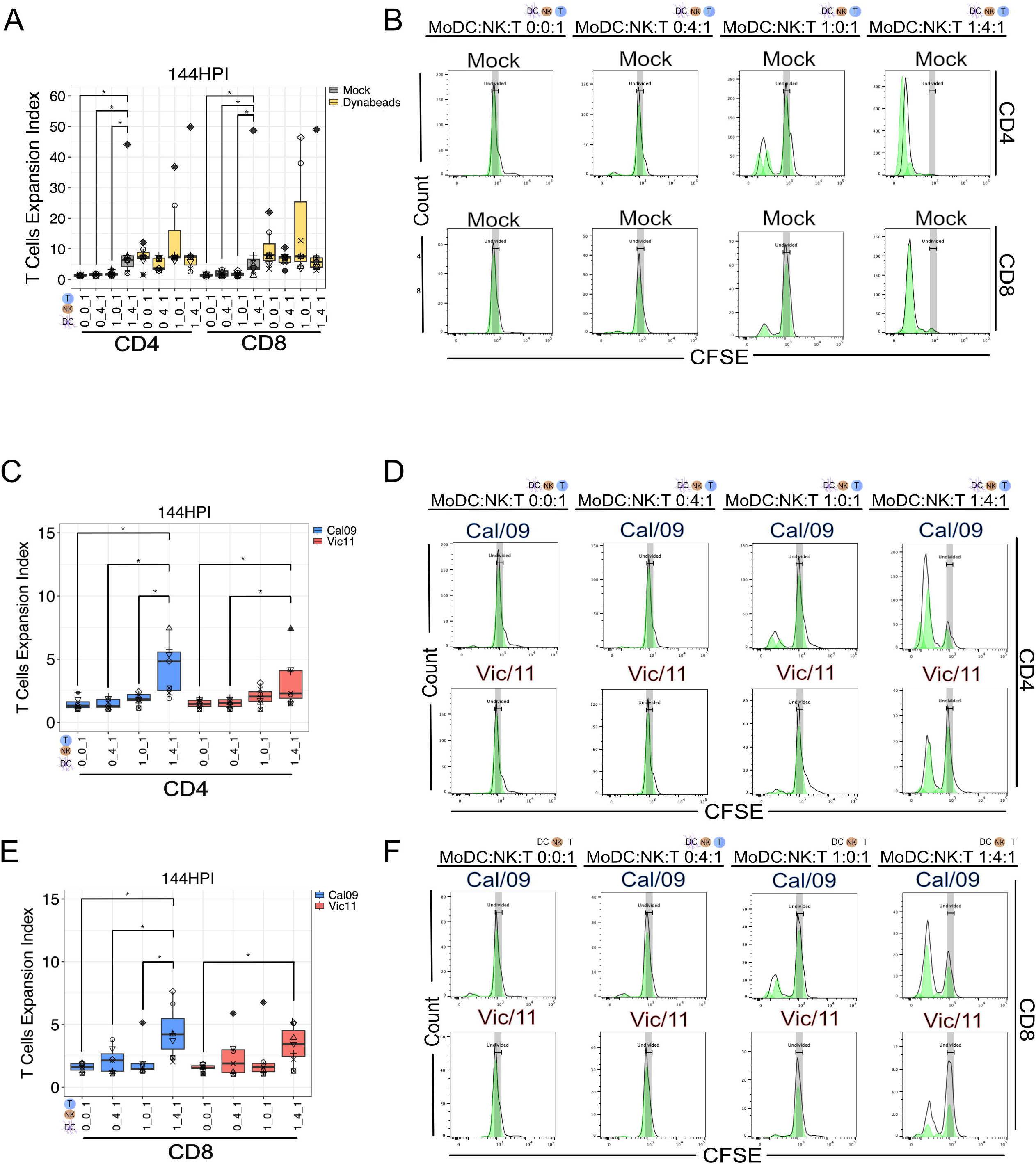
MoDCs trigger expression of proliferation of CD4^+^ and CD8^+^ T cells in a NK cell-dependent manner. **(A)** Summary plot of CFSE-labeled CD4^+^ and CD8^+^ T cell subset expansion index after 143 h co-culture with mock-treated MoDCs and NK cells at MoDC:NK:T cell ratios of 0:0:1, 0:4:1, 1:0:1, 1:4:1 (*n* = 7). Anti-CD3/28-coated Dynabead^®^ treatment at a 1:1 bead-to-cell ratio for the 144 h of the co-culture served as a positive control. **(B)** Representative flow plots of CD4^+^ (top panel) and CD8^+^ (bottom panel) T cell subset expansion index after 143 h co-culture with mock-treated MoDCs and NK cells at MoDC:NK:T cell ratios of 0:0:1, 0:4:1, 1:0:1, 1:4:1. Summary plot **(C)** and representative plots **(D)** of CFSE labeled CD4^+^ T cell subset expansion index after 143 h co-culture with Cal/09 or Vic/11-exposed MoDCs (MOI = 3) and NK cells at MoDC:NK:T cell ratios of 0:0:1, 0:4:1, 1:0:1, 1:4:1 (*n* = 7). Summary plot **(E)** and representative plots **(F)** of CFSE labeled CD8^+^ T cell subset expansion index after 143 h co-culture with Cal/09 or Vic/11-exposed MoDCs (MOI = 3) and NK cells at MoDC:NK:T cell ratios of 0:0:1, 0:4:1, 1:0:1, 1:4:1 (*n* = 7). The data point with the same shape between conditions indicates that it is derived from the same donor. *p < 0.05, Wilcoxon signed-rank test.

### IAV impairs MoDC-NK cell-mediated increase in CD4^+^ and CD8^+^ T cell proliferation in a replication-dependent manner

We then directly compared the effect of IAV on T cell subset proliferation when in triple co-culture (1:4:1). We found that the expansion index for CD4^+^ T cells decreased when exposed to Cal/09 when compared to the mock-treated condition **(Figure 6A-B)**. We also found a significant drop in the expansion index of CD8^+^ T cells when they were exposed to each IAV strain, with Cal/09 inducing a greater expansion index than Vic/11 **(Figure 6A-B)**. Further, we found that CD4^+^ T cells experienced an increase in proliferation when exposed to UV-inactivated Cal/09 when compared to its replication-competent counterpart **(Figure 6C-D)**. Lastly, there was a significant increase in proliferation when exposed to UV-inactivated Cal/09 when compared to UV-inactivated Vic/11 for CD8^+^ T cells **(Figure 6C-D)**.

**Figure 6.**
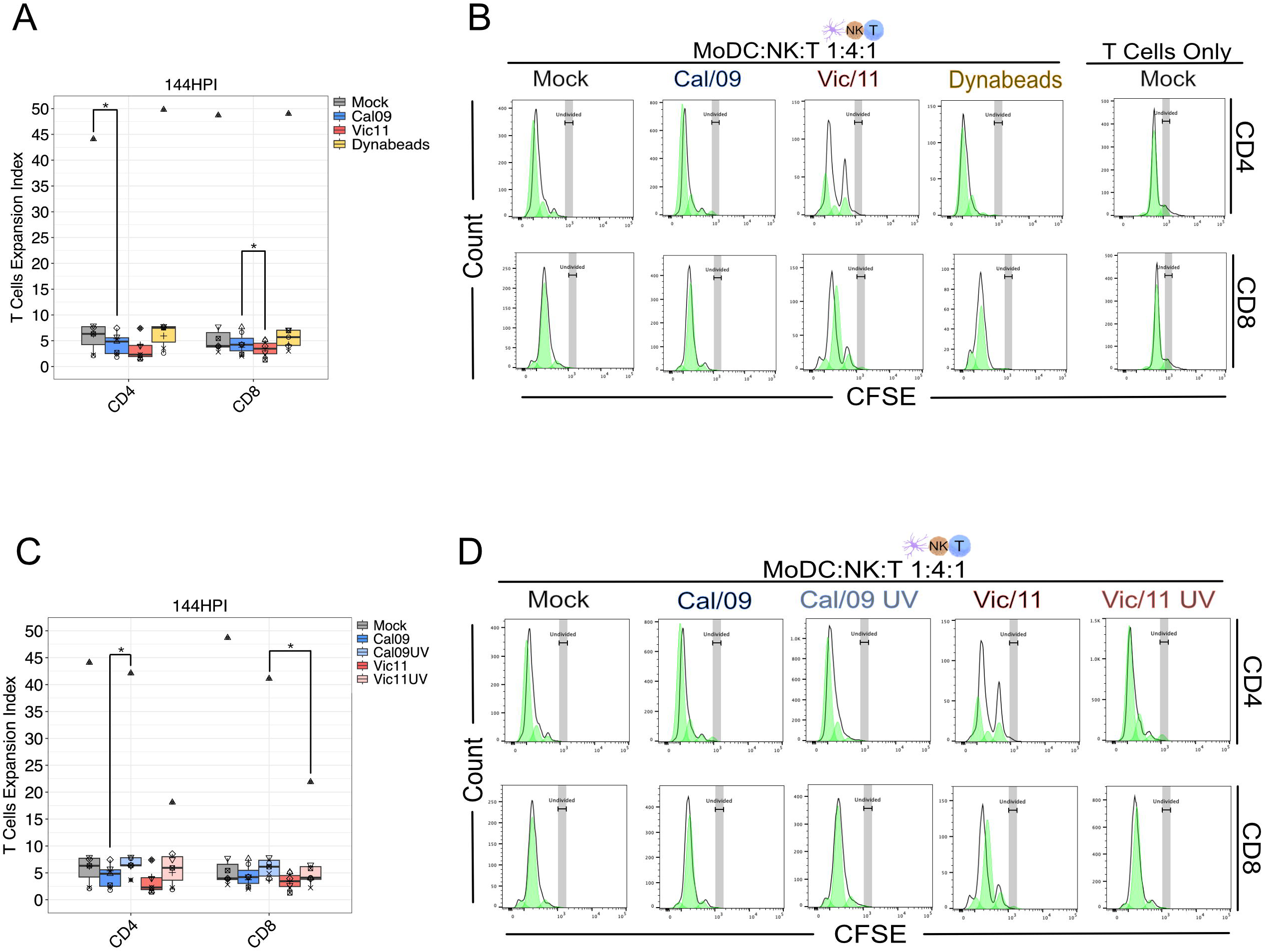
Replication of IAV is required for proliferation of CD4^+^ and CD8^+^ T cells after MoDC-NK cell co-culture. **(A)** Summary plot of CFSE labeled CD4^+^ and CD8^+^ T cell subset expansion index after 143 h co-culture with mock-treated or Cal/09- or Vic/11-exposed MoDCs (MOI = 3) and NK cells at MoDC:NK:T cell ratio of 1:4:1 (*n* = 7). **(B)** Representative flow plots of CFSE labeled CD4^+^ and CD8^+^ T cell subset expansion index after 143 h co-culture with mock-treated or Cal/09- or Vic/11-exposed MoDCs (MOI = 3) and NK cells at MoDC:NK:T cell ratio of 1:4:1 (*n* = 7). **(C)** Summary plot of CFSE labeled CD4^+^ and CD8^+^ T cell subset after 143 h co-culture with mock-treated, replication-competent or UV-inactivated Cal/09 and Vic/11 (MOI = 3) and NK cells at MoDC:NK:T cell ratio of 1:4:1 (*n* = 7). **(D)** Representative flow plots of CFSE labeled CD4^+^ (top panel) and CD8^+^ (bottom panel) T cell subset expansion index after 143 h co-culture with mock-treated, replication-competent or UV-inactivated Cal/09- and Vic/11-exposed MoDCs (MOI = 3) and NK cells at MoDC:NK:T cell ratio of 1:4:1 (*n* = 7). Anti-CD3/28-coated Dynabead^®^ treatment at a 1:1 bead-to-cell ratio for the 144 h of the co-culture served as a positive control. The data point with the same shape between conditions indicates that it is derived from the same donor. *p < 0.05, Wilcoxon signed-rank test.

## Discussion

In this study, we investigated whether influenza A viruses govern crosstalk between human DCs and NK cells and consequently, an impact on naïve T cell activation. We first sought to establish the impact of DC-NK cell crosstalk on the activation of CD4^+^ and CD8^+^ T cell subsets under steady-state, uninfected conditions. We found that crosstalk between human DCs and NK cells leads to activation of naive helper CD4^+^ and cytotoxic CD8^+^ T cells, including upregulation of early (CD69) and late (CD25) markers of surface activation. Further, CD4^+^ and CD8^+^ T cells underwent additional cycles of cell division after exposure to DCs and NK cells, compared to exposure to DCs alone, NK cells alone or in isolation. Consistent with this finding, previous work found that human, resting NK cells become activated and proliferate upon direct interaction with DCs, and these NK cells secrete IFN-γ and acquire cytolytic activity. In turn, these activated NK cells then selectively kill DCs expressing low levels of HLA class I molecules [34]. Similar findings have been reported in mice, whereby DCs activated resting NK cells, including triggering cytolytic activity and IFN-γ production during *ex vivo* culture [35]. Indeed, in our prior study we detected higher levels of IFN-γ in the triple culture conditions, than when T cells were cultured in isolation or with only DCs or only NK cells [25]. These findings provide a potential explanation for our finding of heightened T cell activation, in the absence of viral infection, when cultured with DCs and NK cells. While significant work has characterized DC-NK cell crosstalk, and the role of DC in the activation of naive T cells, our prior study was the first to report how the reciprocal interactions of these innate immune cells positively impact T cell activation and this activation is curtailed by IAV [25]. Here, we have significantly expanded upon that finding to show that both CD4^+^ and CD8^+^ T cell subsets are subject to this activation and IAV-mediated curtailment and further, the pandemic H1N1 Cal/09 also reduced expansion specifically of CD4^+^ T cells.

Several lines of evidence support our observation that IAV regulates DC-NK cell crosstalk leading to impaired T cell activation under *ex vivo* conditions (**Figure 2A**, **4A and 6A**). T cells generally have low basal expression of CD69 than can be rapidly induced by stimulation of the TCR/CD3 complex [37]. Upon exposure of DCs to Cal/09 or Vic/11, the frequency of CD4^+^CD69^+^ and CD8^+^CD69^+^ T cells was significantly lower compared to mock-treated MoDCs. These data are consistent with IAV exposure of MoDCs impairing the ability of T cells to induce CD69 expression on T cells. Notably, CD69 is a hallmark marker of tissue-resident memory T cells, which have been found to play a critical role in conferring heterosubtypic immunity during IAV infection [38–40]. A similar pattern was observed for CD4^+^CD25^+^ and CD8^+^CD25^+^ T cells, with lower frequencies of both subsets detected when MoDCs were exposed to Cal/09 and a lower frequency of CD8^+^CD25^+^ when MoDCs were exposed to Vic/11, compared with mock treatment. CD25 is a high-affinity subunit of the IL-2 receptor and the absence of this subunit could lead to decreased pathogen-specific T cell activation and expansion, further decreasing the development of memory CD8^+^ T cells, and thus protection against future influenza-related disease [41,42].

Consistent with lower levels of the IL-2 receptor, CD4^+^ T cells failed to expand to the same extent when cultured with Cal/09-exposed MoDCs compared to mock treatment.

We reveal a key mechanism underlying the phenotype of impaired T cell activation involves the ability of the virus to undergo replication, because when the Cal/09 was rendered inactivated– without intact replication machinery–T cell activation by DC-NK crosstalk was restored. Intriguing, this did not hold for all IAV strains, as inactivation of the Vic/11 strain failed to restore T cell activation to the same extent as the Cal/09 virus, with the expression of both CD69 and CD25 being significantly higher on both helper CD4^+^ and cytotoxic CD8^+^ T cells when DCs were exposed to Cal/09, compared with Vic/11 (**Figure 2C and 3C**). This raises the possibility IAV strains may be differentially sensed by innate immune pattern recognition receptors. Viral surface glycoproteins, double-stranded RNA, and intracellular viral proteins all have the capacity to activate signal transduction pathways leading to the expression of cytokines with the ability to influence DC-NK cell crosstalk and subsequent T cell activation [43,44]. Significant work has shown that genetically distinct influenza viruses induce pro-inflammatory cytokines to varying levels, with consequences for viral pathogenesis [45–51]. For example, an A/WSN/33 reassortant influenza virus containing NS1–which inhibits NF-κB, IRF3, and IRF7 activation–from the 1918 pandemic virus was more efficient at blocking IFN-regulated genes than the wild-type parental A/WSN/33 [52–54]. However, parsing out whether this difference in cytokine induction is due to differences in viral protein functions, or differential sensing of viral components remains not fully understood. Here we provide evidence that IAV strains may be differentially sensed by pattern recognition receptors, independent of viral replication, with implications for innate immune crosstalk and subsequent induction of adaptive immunity **(Figures 2C & 4C)**.

This study has several limitations. First, our findings were generated using primary human cells cultured *ex vivo* at one cell ratio. This permits investigation of the unique characteristics possessed by human IAV strains, without being adapted by serial passage to infect laboratory mice, however, the impact of these findings on clinical disease outcomes cannot be established. These findings could potentially inform studies in the humanized mouse model to investigate the impact of selective NK cell deletion on DC-mediated priming of naive T cells post-vaccination with inactivated influenza vaccines generated from the pandemic A/California/07/2009 (H1N1) and A/Victoria/361/2011 strains. Second, the influenza infection and vaccination history of the blood donors in this study were unknown. To mitigate this confounder, we isolated the naïve T cell population by negative selection to remove the memory T cell population. This methodology is not 100% efficient and therefore virus-specific memory T cells may be present at low numbers in these co-cultures, which complicates direct comparisons of results between blood donors.

These findings also give rise to the question of which viral protein of the ∼10 encoded by the Cal/09 genome (PB2, PB1, PA, HA, NP, NA, M1, M2, NS1, NS2) is responsible for the impaired T cell activation. This question has direct implications for influenza mRNA-based vaccine design, as genetically separating the viral protein that elicits a protective immune response, including harboring conserved T cell epitopes, may be useful in the selection of the vaccine antigen candidate most effective at eliciting a robust T cell response. This could be performed by generating mRNA constructs encoding each viral protein followed by *in vitro* transcription and delivery into DCs individually. Determining which viral protein, with peptide presented and cross-presented by HLA class I and II MoDCs triggers the most effective antigen-specific T cell response will be an important future endeavor.

## Acknowledgments

We would like to thank the study participants and Drs. Ellermeier and Samanta for careful reading of the manuscript.

## Author Contributions

Conceptualization: L.M.K and B.S.L. Experiments: C.M and C.U. Analyses: L.M.K. Writing:

L.M.K. Funding: L.M.K and B.S.L.

## Nomenclature

Dendritic cells (DC); Natural Killer cells (NK cells); monocyte-derived dendritic cell (MoDC); Influenza A Virus (IAV), Peripheral blood mononuclear cells (PBMCs).

---

**Figure S1. Lineage gating schematic representing CD69^+^ and CD25^+^ CD4^+^ and CD8^+^ T cells.** Diagram of lineage gating tree is used to identify **(A)** CD4^+^CD69^+^ T cells, **(B)** CD8^+^CD69^+^ T cells, **(C)** CD4^+^CD25^+^ T cells and **(D)** CD8^+^CD25^+^ T cells gated using an FMO control to inform gating location. Schematic of lineage gating tree and proliferation modeling histogram to identify **(E)** proliferating CD4^+^ T cells and **(F)** proliferating CD8^+^ T cells by flow cytometry.

