## Supplementary figures and images for "Influenza A Virus Disruption of Dendritic Cell-Natural Killer Cell Crosstalk Impacts Activation of Helper and Cytotoxic T cell Subsets"

### Figure S1

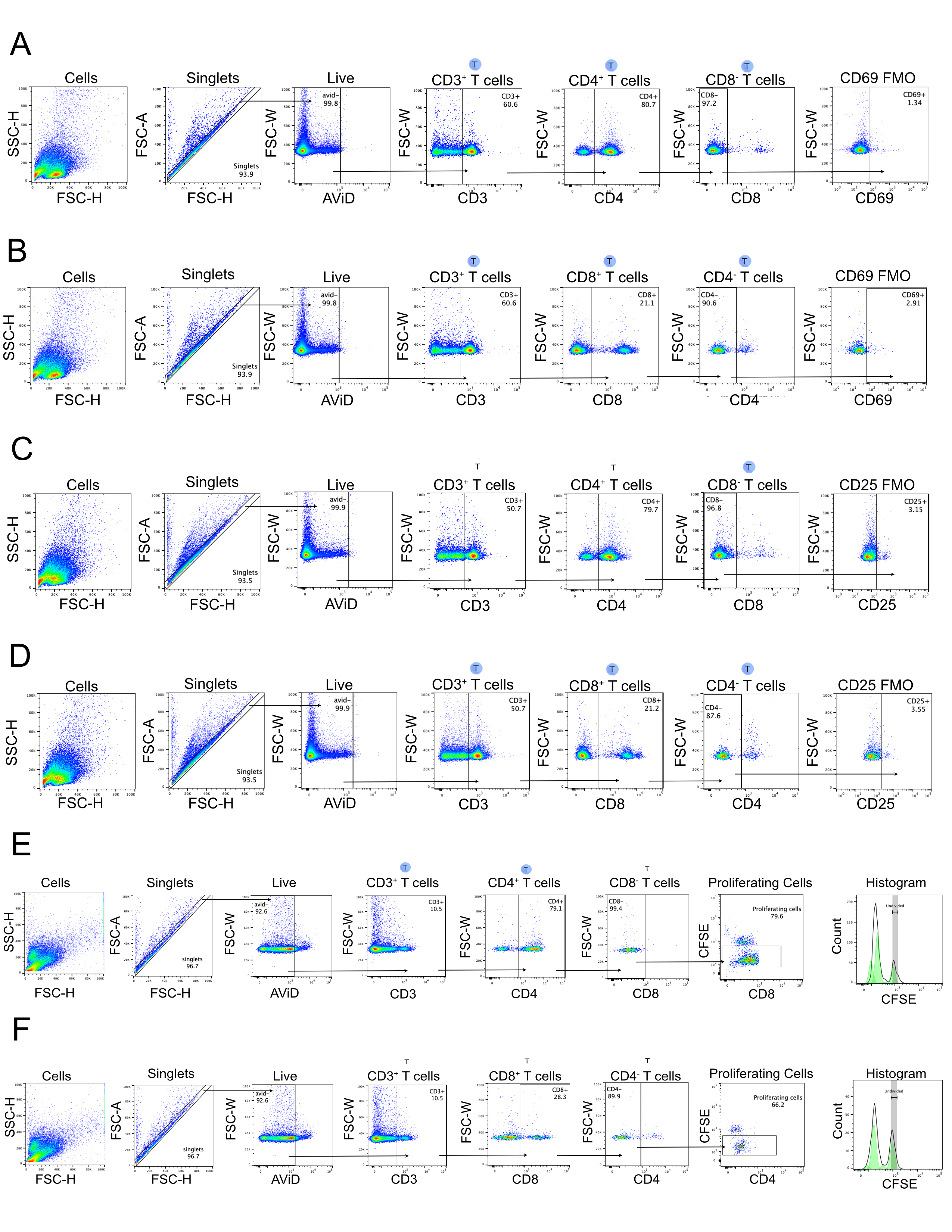
